# Integrating Optogenetic Stimulation of Olfactory Bulb Glomeruli with Foot Shock Fear Conditioning: A Robust Method for Investigating Olfactory-based Fear Conditioning

**DOI:** 10.1101/2024.03.25.586672

**Authors:** Praveen Kuruppath

**Affiliations:** University of Colorado Anschutz Medical Campus, Aurora, Colorado 80045, USA

**Keywords:** Optogenetics, channelrhodopsin-2, foot shock, light zone, olfactory bulb, safe zone

## Abstract

The integration of optogenetic techniques with traditional behavioral paradigms has provided novel insights into the neural mechanisms underlying olfactory-based fear conditioning. Olfactory cues are potent triggers for fear responses, and understanding the intricate neural dynamics involved in olfactory fear learning is crucial for unraveling the complexities of aversive memory formation. In this study, a robust method is presented that combines optogenetic stimulation of olfactory bulb glomeruli with foot shock fear conditioning to investigate olfactory-based fear learning in mice. By merging optogenetic manipulation with behavioral assays, a comprehensive framework for studying the mechanisms of olfactory fear conditioning is provided. This method offers new avenues for exploring the neural dynamics of adaptive responses to olfactory threats and may have implications for understanding fear-related disorders.

## Introduction

The ability to perceive and respond to environmental stimuli are a fundamental survival mechanism in both humans and animals. Olfactory cues play a pivotal role in this process serving as a powerful trigger for fear response. The study of olfactory-based fear conditioning provides valuable insights into the neural mechanisms underlying the formation and consolidation of aversive memories (Meissner-Bernard et al., 2019; Mouly and Sullivan, 2009).

Understanding the intricate mechanisms underlying the formation and consolidation of fear-associated memories is an important goal in neuroscience research (Meissner-Bernard et al., 2019). Among the sensory modalities, olfaction holds a unique position, as it is known to elicit a profound and long-lasting emotional response. The olfactory system’s ability to swiftly and powerfully influence the formation of fear memories has made it a subject of great interest in the field of behavioral neuroscience (Silvas-Baltazar et al., 2023).

To explore this process in greater detail, researchers have used cutting-edge techniques that allows precise control of neural circuits. One such technique, Optogenetics has revolutionized the field of neuroscience by enabling the manipulation of specific neuronal populations with exceptional precision (Kushibiki et al., 2014; Pastrana, 2011). Combining optogenetics with traditional behavioral paradigms, such as fear conditioning provides a powerful platform for investigating the neuronal underpinnings of olfactory-based fear learning.

In this study, a robust method is presented that integrates optogenetic stimulation of olfactory bulb glomeruli with foot shock fear conditioning. This approach offers a unique opportunity to dissect the intricate neural pathways responsible for encoding and modulating olfactory-associated fear memories. By merging the sophistication of optogenetics with the well-established principles of fear conditioning, this method opens new avenues for exploring the neural dynamics that drive adaptive behavioral responses to olfactory threats.

## Materials and Methods

### Experimental animals

All animal procedures conformed to National Institutes of Health guidelines. Mice were bred in-house and were maintained on a 12 h light/dark cycle with food and water ad libitum. Experimental animals were generated by crossing heterozygous tetO-ChIEF-Citrine (tetO-ChIEF-Citrine+/−) with homozygous OMP-tTA mice (OMP-tTA-/-). The offspring were OMP-tTA+/-/ tetO-ChIEF-Citrine+/- (OMP-ChIEF) (Cheetham et al., 2016; Kuruppath et al., 2021; Kuruppath and Belluscio, 2021).

### Genotyping

OMP-ChIEF pups were identified by the visualization of fluorescence in the nose and OB of P0-P2 pups under epifluorescence illumination.

### Animal preparations

#### Construction of implantable optical fibers

Implantable optical fibers and patch cables were prepared as described previously (Sparta et al., 2012a; Ung and Arenkiel, 2012). Briefly, Prepare the heat-curable fiber optic epoxy by combining 100 mg of hardener with 1 g of resin. Next, use a wedge-tip carbide scribe to measure and score approximately 35 mm of 125 μm fiber optic containing a 100 μm core. Ensure to avoid cutting the fiber entirely to prevent core damage. Insert an LC ceramic ferrule with a 127 μm bore into a vice, ensuring the convex side faces downward. Carefully insert the fiber optic into the ferrule, ensuring smooth insertion and a slight protrusion from the convex end. Apply a single drop of epoxy to the flat end and heat it with a heat gun until it turns black, ensuring the epoxy fills the ferrule before curing. Remove any excess epoxy from the ferrule’s sides to prevent obstruction during interfacing. Polish the ferrule’s convex end using a fiber optic polishing disc on aluminum oxide polishing sheets of different grades placed on the polishing pad. Use circular rotation patterns and polish in four grades: 6, 3, 1, and 0.2 μm grit. Cut the fiber optic at the flat end to the desired length. Test the implant by connecting it to the laser using the coupler cord. Ensure the polished implant end makes direct contact with the opposing ferrule and maintains a consistent light output measured at the implant fiber tip (approximately 20mW – 22mW). Implants with weak focal points near the fiber optic tip indicate poor quality. Finally, store the completed fiber implants in foam until ready for use.

#### Construction of patch cables

Prepare the heat-curable fiber optic epoxy by combining 100 mg of hardener with 1 g of resin as mentioned above. Then, measure and score an appropriate length of 220 μm fiber optic with a 200 μm core using a wedge-tip carbide scribe. Insert the fiber optic into a piece of furcation tubing slightly longer than the fiber, with an inner diameter slightly larger than the fiber’s width. Strip approximately 25mm from one end of the fiber and insert it firmly into the metal end of a Multimode FC Ferrule assembly with a 230 μm bore, ensuring it protrudes from the ferrule end. Secure the connection with cyanoacrylate at the metal end, then cover it with a Connector Boot and polish the ferrule end using a fiber optic polishing disc on aluminum oxide polishing sheets in four grades: 6, 3, 1, and 0.2 μm grit. Strip and insert the other end of the fiber into a LC ceramic ferrule (230 μm i.d. boreti with the convex end distal. Apply epoxy to the flat end and heat it until curedti. Polish the ferrule’s convex end using fiber optic polishing disc on aluminum oxide polishing sheets. Check for any excess epoxy or cracks to the fiber core. The fiber core should look like a black concentric circle. If there is excess epoxy on the core, repolish with the lowest grades of polishing paper.

Slide a LC ferrule sleeve over the convex end until the midpoint. Cover the furcation tubing and sleeve with heat-shrink tubing, securing and protecting the connection through heating. Finally, test the coupler’s performance by connecting it to the laser source and measuring the light output through an optical power meter, ensuring light loss doesn’t exceed 30% from the laser output to the coupler output.

#### Surgical implantation

To begin the surgical procedure, mice were anesthetized via intraperitoneal injection of a Ketamine/Xylazine mixture at doses of 100 and 10 mg/kg body weight respectively. Each animal was then securely positioned on a stereotaxic frame, and their head was immobilized with bar ties attached to each temporal side of the skull. Hand warmers were used to maintain optimal body temperature throughout the procedure (Grabber, Grand Rapids, MI, USA).

Once the animals exhibited no response to foot pinching, indicating successful anesthesia, a craniotomy was performed above each olfactory bulb (OB), precisely 4.3 mm anterior from bregma. Fiber optic pins were then implanted on the dorsal surface of each OB (Figure 1*a*), as previously described (Li et al., 2014; Sparta et al., 2012; Ung and Arenkiel, 2012). Post-surgery, mice received a Ketoprofen injection (5 mg/kg) for pain management. Subsequently, animals were allowed a one-week recovery period in their home cage.

**Figure 1.**
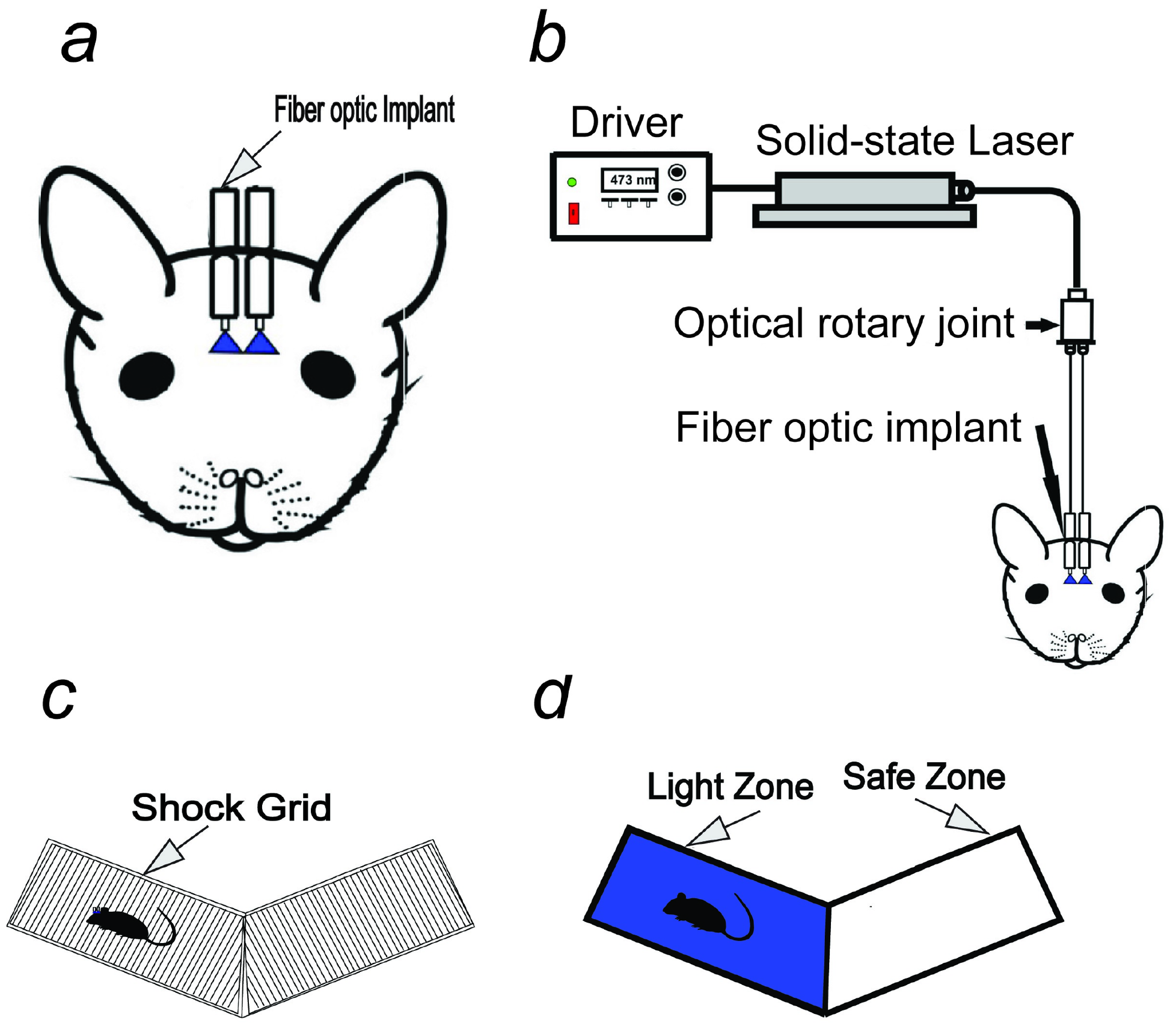
Optogenetic stimulation of olfactory bulb. ***a***, Schematic diagram of the fiber implant. ***b***, Behavioral setup. ***c***, “V” maze paved with electric shock grid for behavior training. ***d***, “v’ maze without electric shock grid for behavior testing. Light zone indicates the area where the light stimulation is delivered. Safe zone indicates the area where mice can escape from the light stimulation and foot shock.

#### Optical activation paired with avoidance conditioning using foot shock

After the animals recovered from surgery (one-week post-surgery), behavioral training commenced using a modified ‘two-arms maze,’ derived from the ‘Y’ maze design. This maze comprised two equally sized open arms. Each open arm was fitted with an electric grid shock floor. The mice were connected to a 400 μm core-diameter optical fiber linked to a 473 nm solid-state variable-power laser (LaserGlow Technologies, Toronto, Canada, Figure 1*b*) and allowed a 15-minute habituation in the maze (Figure 1*c*, *Movie1)*. Their exploration time in each arm was recorded to identify arm preference. The arm favored by the mice was designated as the Light zone, while the opposite arm became the Safe zone (Figure 1*d*). In cases where no clear preference was evident, the Light zone was randomly assigned.

After habituation, mice underwent daily ten-minute maze training sessions, occurring sixty minutes before testing to enhance learning. During training, light stimulation paired with mild foot shocks was applied as the mice fully entered the Light zone (Figure 1*d*). The Safe zone remained accessible to the mice to avoid the foot shocks. The light stimulation involved ten 50 ms pulses with 150 ms intervals, triggered externally by a Master-8 timer (A.M.P.I, Jerusalem, Israel) (Li et al., 2014; Erofeev et al., 2019; Quinlan et al., 2019). Light power output was set between 20-22 mW based on prior studies (Lin, 2011a, 2011b, 2012; Sparta et al., 2012b).

By using a stand-alone shock generator (Med Associates, USA), a mild 0.65 mAmps, 5-second-long foot shock was delivered 2 -second after the light stimulation, synced with the Master-8 timer. The mice were conditioned to move to the Safe zone upon receiving the light stimuli and foot shock in the Light zone.

#### Light zone aversion assessment task

The Light Zone aversion assessment test lasted for two days and included reinforcement training sessions prior to each day’s testing phase. Testing commenced 60 minutes post-reinforcement training. To ensure unbiased testing conditions, the electric grid shock floor was removed from the “two-arms maze,” eliminating foot shock exposure during the testing sessions.

Mouse activity within the maze was evaluated in blocks of three trials, each lasting 15 minutes. Initially, mice explored the arena without light stimulation to establish baseline behavior. Subsequent trials focused on assessing arm preferences post-foot shock training, specifically targeting the Light Zone and Safe Zone as previously designated areas.

Movement within the “two-arms maze” was tracked using Any-maze video tracking software (Stoelting, IL, USA) to calculate time spent in each arm. Additionally, heatmaps were generated for every trial, visually displaying the duration spent by mice in distinct areas of the arena. Color ranges within the heatmap represented varying durations, with blue indicating the shortest and red indicating the longest time spent in specific regions.

## Results

Experience significantly influences odor perception and behavioral responses, with learned odor-context associations often persisting throughout an animal’s life. Yet, the precise cellular and neural circuit mechanisms responsible for olfactory learning and memory remain poorly understood.

Here, foot shock avoidance training was performed paired with either left or right OB stimulation. After training, the Light zone avoidance response to OB stimulation was tested. Initially, as a control, the time spent in each arm was calculated by allowing the mice to freely explore the arena in the absence of stimulation (Movie2). The purpose of the baseline behavior analysis was to determine whether the mice had any preference for a particular arm of the “two-arms maze” after the foot shock training. Baseline data showed that mice spent equal amounts of time in both arms of the maze (Left zone – 443 ± 14.08 s, Right zone-457 ± 14.08 s, P = 0.8179, Figure 2a). Then, light stimulation was delivered to the unilateral OB. It was found that mice avoided the Light zone during unilateral OB stimulation and spent most of their time in the Safe zone (Light zone - 82.25 ± 27.01 s, Safe zone - 817.75 ± 27.01 s, P = <0.0001, Figure 2b, 2c, Movie 3, n = 4). Additionally, testing was conducted to determine whether the avoidance response to the light stimulation was equally probable in both arms of the maze by delivering the stimulation when the mice reached the Safe zone. It was found that mice avoided the Safe zone during the light stimulation, indicating that the avoidance response is clearly linked to the olfactory information, but not to spatial information.

**Figure 2.**
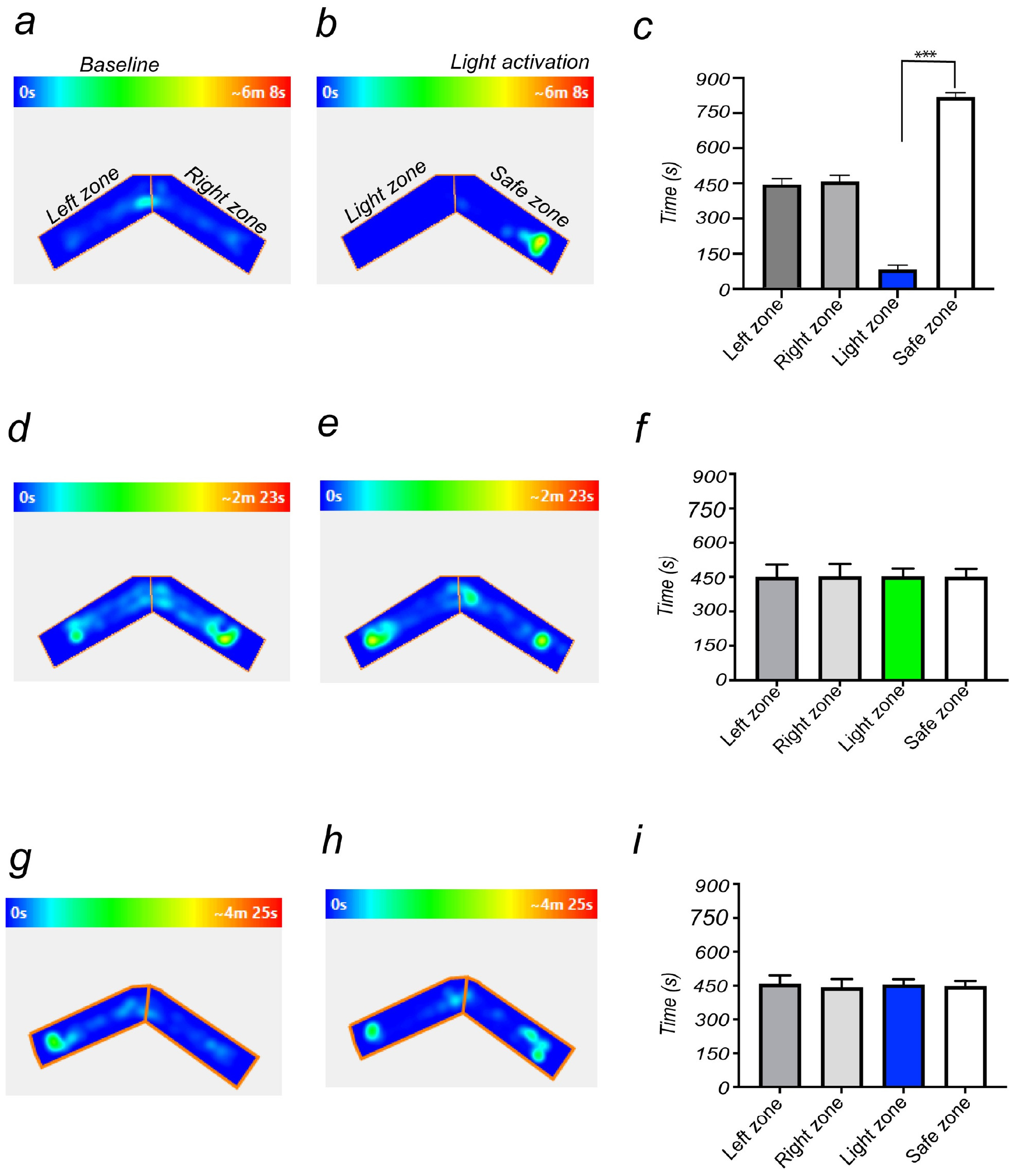
Light zone avoidance response to OB stimulation. ***a***, *b*, an example of heat-map showing animal’s position in “two-arms maze”, during baseline (*a*) and OB stimulation (*b*). *c*, Average amount of time explored in each zone in baseline (LeI zone, Right zone) and OB stimulation trials (Light zone, Safe Zone). ***d, e***, Heat-map of mouse position during baseline (***d***) and green light stimulation (***e***). ***f***, Average amount of time spent in each zone in green light stimulation. ***g, h***, an example of heat-map showing WT animal’s position in “two-arms maze”, during baseline (***g***) and OB stimulation (***h***). ***i***, Average amount of time explored in each zone in baseline (LeI zone, Right zone) and OB stimulation trials (Light zone, Safe Zone).

To verify that the observed responses from the light stimulation were the result of activation of the ChIEF-expressing neurons and not from the use of light as a visual cue, green light (540nm, output power, 20-22mW), which does not activate ChIEF, was used. Mice are much less sensitive to such long wavelength light (Denman et al., 2017, 2018; Peirson et al., 2018). During the green light stimulation, significant behavioral differences from the baseline behavior were not observed (Left zone - 459 ± 36.13 s, Right zone - 441 ± 36.13 s, P = 0.81, Light zone - 441 ± 23.88s, Safe zone - 459 ± 23.88 s, P = 0.72, Figure 2d-f), confirming that the mice did not use visual cues to perform the task. The test was also conducted with wild-type mice, confirming that visual cues were not involved in solving the behavioral task (Left zone - 457.8 ± 40.21 s, Right zone - 442.3 ± 40.21 s, P = 0.86, Light zone - 451 ± 18.03 s, Safe zone - 449 ± 18.03 s, P = 0.96, Figure 2g-i, n = 4). These results indicate that OMP-ChIEF mice were using only the light activation of the olfactory system as the cue to solve the behavioral task, rather than light detection via other modalities.

The findings indicate that linking optogenetic stimulation with foot shock fear conditioning triggers a strong avoidance response that is measurable. This approach appears to facilitate the integration of distinct ontogenetically induced olfactory information into fear memory, which can be quantified behaviorally.

## Discussion

The present study introduces a robust method for investigating olfactory-based fear conditioning by integrating optogenetic stimulation of olfactory bulb glomeruli with foot shock fear conditioning. The observed avoidance behavior in response to the combined optogenetic and foot shock stimuli demonstrates the effectiveness of the proposed methodology. In the experiment, the mice exhibited a clear preference for the Safe zone, indicating a learned association between the optogenetic activation of olfactory bulb glomeruli and aversive foot shock. Importantly, the specificity of this response to olfactory information rather than spatial cues was confirmed through control experiments using green light and wild-type mice. These control experiments strengthen the validity of the results, suggesting that the observed behavioral changes are indeed linked to the optogenetic manipulation of the olfactory system.

The integration of optogenetics with traditional fear conditioning paradigms offers a powerful platform for studying the neural dynamics underlying olfactory-based fear learning. The ability to precisely control specific neuronal populations using optogenetics provides a level of precision that enhances the mechanistic understanding of fear memory formation. Moreover, the use of foot shock as an aversive stimulus in conjunction with olfactory stimulation mimics a real-world scenario, aligning with the ecological relevance of fear responses to olfactory threats. The experimental design, including the use of a modified ‘two-arms maze’ and the Light Zone aversion assessment task, provides a comprehensive behavioral framework for evaluating olfactory fear responses.

Despite the promising results, it is essential to acknowledge certain limitations. For instance, the specific mechanisms by which olfactory information is integrated into fear memory need further exploration. Future studies could investigate the role of specific neural circuits and molecular pathways involved in this process, potentially using additional techniques such as electrophysiology and molecular biology assays.

In conclusion, the presented methodology successfully integrates optogenetic stimulation with foot shock fear conditioning, providing a robust approach for investigating olfactory-based fear learning. The present method contributes to the growing body of knowledge on the neural mechanisms underlying olfactory related fear memory and open new avenues for research in the field of behavioral neuroscience. Further exploration of the molecular and circuit-level details of olfactory fear conditioning will deepen our understanding of adaptive responses to olfactory threats and may have implications for the development of therapeutic interventions for fear-related disorders.

## Supporting information

Habituation

Baseline Behavior

Light zone avoidance Behavior

## Acknowledgement

This work was performed at the National Institute for Neurodegenerative Disorders and Stroke, and it was supported by the National Institutes of Health intramural program.

## Additional Information

The author declare no competing interests.

## Notes

### Competing Interest Statement

The authors have declared no competing interest.

